# Lattice micropatterning for cryo-electron tomography studies of cell-cell contacts

**DOI:** 10.1101/2020.08.30.272237

**Authors:** Leeya Engel, Claudia G. Vasquez, Elizabeth A. Montabana, Belle M. Sow, Marcin P. Walkiewicz, William I. Weis, Alexander R. Dunn

## Abstract

Cryo-electron tomography is the highest resolution tool available for structural analysis of macromolecular complexes within their native cellular environment. At present, data acquisition suffers from low throughput, in part due to the low probability of positioning a cell such that the subcellular structure of interest is on a region of the electron microscopy (EM) grid that is suitable for imaging. Here, we photo-micropatterned EM grids to optimally position endothelial cells so as to enable high-throughput imaging of cell-cell contacts. Lattice micropatterned grids increased the average distance between intercellular contacts and the thicker cell nuclei such that the regions of interest were sufficiently thin for direct imaging. We observed a diverse array of membranous and cytoskeletal structures at intercellular contacts, demonstrating the utility of this technique in enhancing the rate of data acquisition for cellular cryo-electron tomography studies.

## Introduction

Cryo-electron tomography (cryo-ET) is a three-dimensional electron microscopy (EM) technique uniquely capable of imaging the interior of frozen cells at the nanometer scale (1–4). While other EM techniques have contributed to our understanding of macromolecular organization in cells, they do not maintain macromolecules in their native state within a hydrated cell. For example, platinum replica electron microscopy is a productive technique for studying cytoskeletal organization, but it strips away the cell membrane and introduces artifacts associated with dehydration and chemical fixation (5). Preservation of the membrane, cytoskeletal structure and associated proteins makes cryo-ET particularly suited to assessing the ultrastructure of cell-cell contacts (6)

Cryo-ET is limited by the throughput of data collection, as it is time consuming to collect a sufficient number of high-quality tilt-series (7). This low throughput has a number of sources, including the mechanical limitations of the stage requiring refocusing, the software operating the microscope, and the ability to locate the region of interest. The latter issue is particularly limiting for *in situ* cryo-ET due to the challenge of identifying target complexes in the complex cellular milieu and the improbability of a cell or a subcellular region of interest being fortuitously positioned in an imageable region of the EM grid. We and others have addressed the latter limitation by applying photo-micropatterning to confine single cells to extracellular matrix (ECM) islands of predefined shapes in the centers of EM grid-squares (8, 9). However, reliably positioning subcellular structures or contacts between cells remains an unmet challenge.

A longstanding challenge has been to visualize the formation and remodeling of cell-cell junctions in endothelial cells (ECs). ECs line the vasculature and actively remodel their intercellular junctions during inflammation, wound healing and vessel remodeling. Despite their critical importance to vascular health, relatively little is known about the earliest and last stages of cell-cell junction formation and dissassembly, as the ultrastructure of remodeling junctions is difficult to study using conventional imaging modalities. Here, we expand on EM grid micropatterning to facilitate cryo-ET of endothelial cell protrusions and intercellular contacts.

To image endothelial junctions by cryo-ET, we photo-micropatterned a continuous ECM lattice aligned to the EM grid that promotes the assembly of thin cell-cell contacts in imageable regions of the grid (Fig. 1). Our quantitative analysis indicates (a) that a latticework of ECM can improve the positioning of intercellular contacts relative to the EM grid and (b)that the lattice micropattern directs the thicker cell nuclei away from the centers of EM grid-squares such that cell-cell contacts can be sufficiently thin for imaging by cryo-ET. Our cryo-tomograms revealed a rich variety of sub-cellular structures in these regions, including contacting filamentous (F)-actin rich membrane protrusions between cells, bundles of intersecting membrane protrusions, and a range of vesicle shapes and sizes within and outside of the plasma membrane. Lattice micropatterning of EM grids can be generalized to facilitate structural studies of a variety of other systems such as neuronal and immunological synapse formation. This study thus advances *in situ* cryo-ET by refining techniques that dramatically increase the throughput of *in situ* cryo-ET data acquisition.

**Fig. 1.**
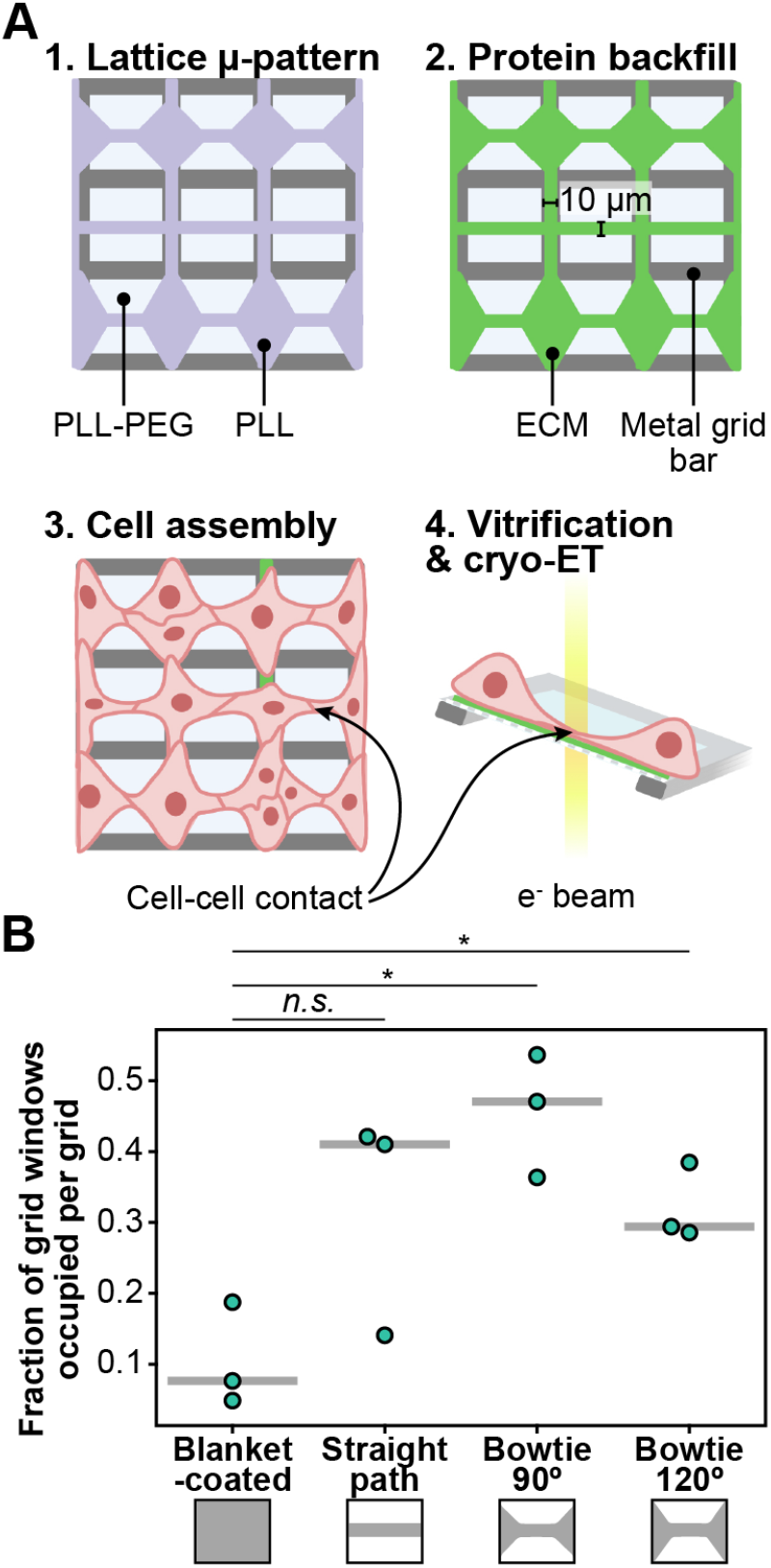
Lattice micropatterning increases the number of endothelial cell-cell contacts located in imageable regions of EM grids. (A) Schematic of lattice micropatterning method. Following a PEG-PLL surface treatment, PEG is selectively degraded by maskless photopatterning (purple). Protein backfill creates an ECM lattice (green) which promotes the assembly of cell-cell contacts in the windows between grid bars. (B) Micropatterning increased the number of optimally positioned cell-cell junctions per EM grid. Each point represents a different EM grid. Median for all measurements indicated by the gray line. N = 3 grids for each ECM condition. 430 measurements of blanket-coated grid-squares, 108 measurements of 90^*o*^ bowtie grid-squares, 108 measurements of 120^*o*^ bowtie grid-squares, and 206 measurements for straight micropatterned grid-squares. Welch’s *t* tests: *P < 0.05; ns = not significant.

## Results and Discussion

To establish an efficient pipeline for imaging endothelial cell-cell contacts by *in situ* cryo-ET, we micropatterned EM grids with a continuous lattice of ECM aligned to the grid bars. Intersections between ECM tracks attracted the thicker cell bodies while thin, contact-forming protrusions extended along the thin, 10 µm wide regions of ECM, where they could be imaged (Fig. 1A). We evaluated the ability of three micropattern variations to direct cell-cell contact assembly. “Straight” micropatterns consisted of 10 µm wide intersecting tracks, while “bowtie” micropatterns consisted of 10 µm tracks that widened into 90^*o*^ or 120^*o*^ diamond shapes at ECM track intersections. We compared the number of cell-cell contacts on imageable regions of EM grids that were blanket-coated with ECM to the number of cell-cell contacts on imageable regions of EM grids that were micropatterned with an ECM lattice (Fig. 1B). Cell-cell contacts, indicated by the presence of vascular endothelial cadherin (VE-cad), were observed on only 10.4 +/- 4.2 (S.E.M.)% of grid-squares on EM grids that were blanket-coated with ECM (Fig. 1B, 2A). In contrast, cell-cell contacts were observed on an average of 32.4 +/- 9.1 (S.E.M.)% of grid-squares for straight ECM micropatterns, 45.7 +/- 5.0 (S.E.M.)% of grid-squares for 90^*o*^ bowtie micropatterns, and 32.1 +/- 3.1 (S.E.M.) for micropatterns with 120^*o*^ bowtie micropatterns. Thus, lattice micropatterning resulted in a significant, greater than threefold increase in the instances of cell-cell contacts in imageable regions of EM grids seeded with ECs.

While previous studies in other contexts have established that cell-cell junction position can be regulated by ECM micropatterning, the nuclei of cell pairs on micropatterned ECM islands tend to be located in close proximity to the cell-cell junctions (10, 11) (Fig. S1). Cell nuclei in endothelial cells are typically several microns in height (Fig. S2). Thus, proximity of the nucleus to the cell-cell contact precludes the possibility of imaging the cell-cell contact using cryo-ET without an additional sample thinning step (Fig. S2-3). While thinning frozen cells by cryo-focused ion beam milling (cryo-FIB) is gaining traction as a preparation step for cellular cryo-ET, it requires specialized machinery that is not yet widely available and can introduce contamination or devitrification to frozen samples (12). Moreover, cryo-FIB restricts experimental throughput due to its slow speed and complexity. Lattice micropatterning improves the likelihood of a well-positioned cell-cell contact falling below the 500 nm thickness threshold for direct imaging (3) by increasing the distance between thick nuclei and intercellular junctions (Fig. 2).

**Fig. 2.**
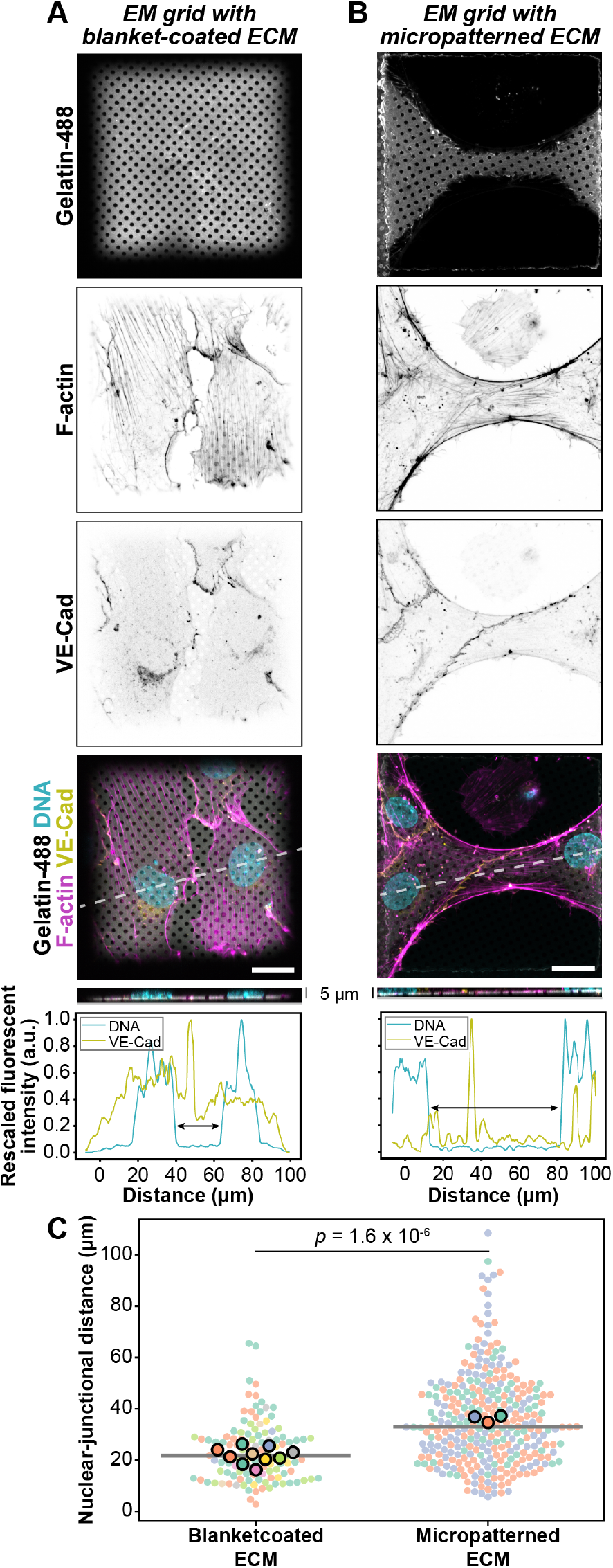
Lattice micropatterning increased the distance between nuclei and cell-cell junctions in ECs cultured on EM grids. (A, B) Representative image of fixed ECs on EM grids blanket-coated (A) and micropatterned (B) with Oregon Green 488-gelatin, and immunostained for F-actin, VE-Cad and DNA. Dashed line indicates position of the Z-stack reslice and line profile of DNA (teal) and VE-cad signals (green). (C) Each point represents an individual measurement of nuclear-junctional distance and each color represents a different EM grid. The mean of the measurements from each grid is indicated by a point outlined in black. Median for all measurements is indicated by the gray line. N = 10 grids, 120 measurements for blanket-coated condition, N = 3 grids, 320 measurements for micropatterned condition. P-value calculated using Welch’s *t* test comparing grid replicates. Scale bars, 20 µm.

We used atomic force microscopy (AFM) to examine the topography and heights of EC cell-cell contacts. While we observed cell-cell contacts that spanned the 10 µm width of the micropattern, consistent with the contacts observed by fluorescence microscopy in figures 3A and S3, we also observed thinner, micron and submicron scale protrusions between cells that appeared to use the ECM micropattern as tracks for contact initiation or retraction (Fig. 3B, S4). We measured endothelial cell-cell contact heights on blanket-coated gelatin and micropatterned glass substrates. Most of the contacts measured were below 500 nm high and we noted an anecdotal relationship between cell-cell contact width and height. Cell-cell contacts below the 500 nm threshold were also below 10 µm wide for both micropatterned and blanket-coated EM grids (Fig. 3C).

**Fig. 3.**
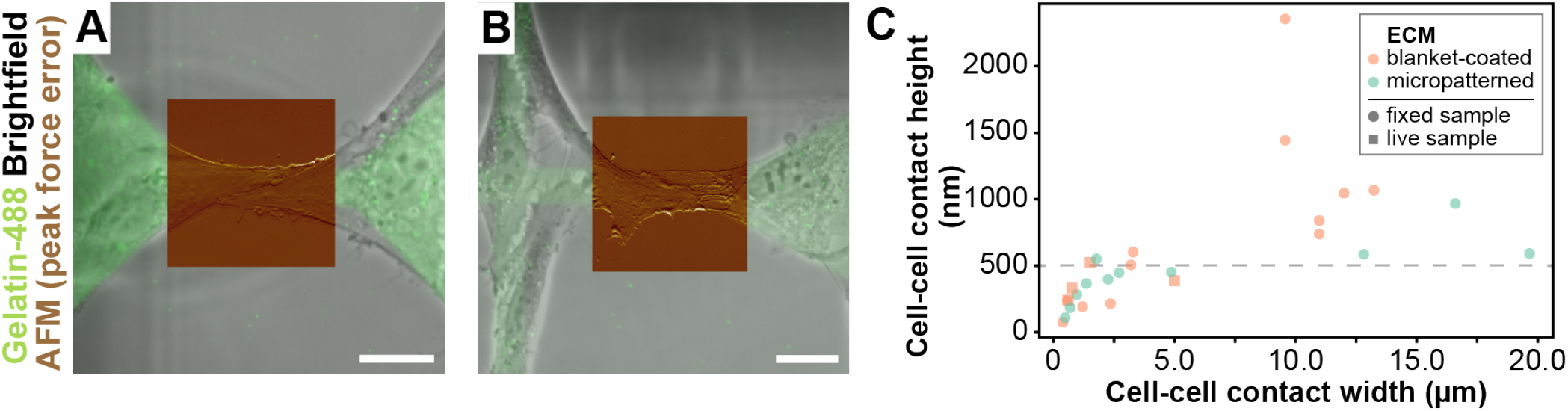
(A,B) Representative brightfield image of ECs on micropatterned glass (grayscale) overlaid with fluorescence image to show micropattern (green). Insets (AFM peak force error) show the topography of continuous (A) and filamentous (B) cell-cell contact phenotypes. Scale bars, 20 µm. (C) Measurements from the AFM height sensor channel show that EC cell-cell contact height can be below 500 nm, the thickness threshold for cryo-EM samples (3). Height is plotted as a function of width for EC cell-cell contacts on blanket-coated (orange) and micropatterned (green) glass samples. N=12 contacts for micropatterned condition. N=15 contacts for blanket-coated condition.

Lattice micropatterned EM grids generated reproducible cellular samples for cryo-ET. The grid atlas in Fig. 4A shows how lattice micropatterning can be used to orient cells (vertically or horizontally) on grid-squares, enabling the programming of visual labels for correlative EM techniques. For example, orienting cells in a single quadrant of the grid perpendicular to the direction of the cells in the other quadrants makes the location of the grid center apparent during TEM, enabling correlation with grid maps generated by light microscopy. In our hands, lattice micropatterning led to a dramatic increase in the ease of data acquisition. Once we had established optimal cell seeding density and freezing parameters, we found that we did not need to screen vitrified grids prior to imaging, as is typically required in order to identify a sufficient number of cellular subregions suitable for imaging.

**Fig. 4.**
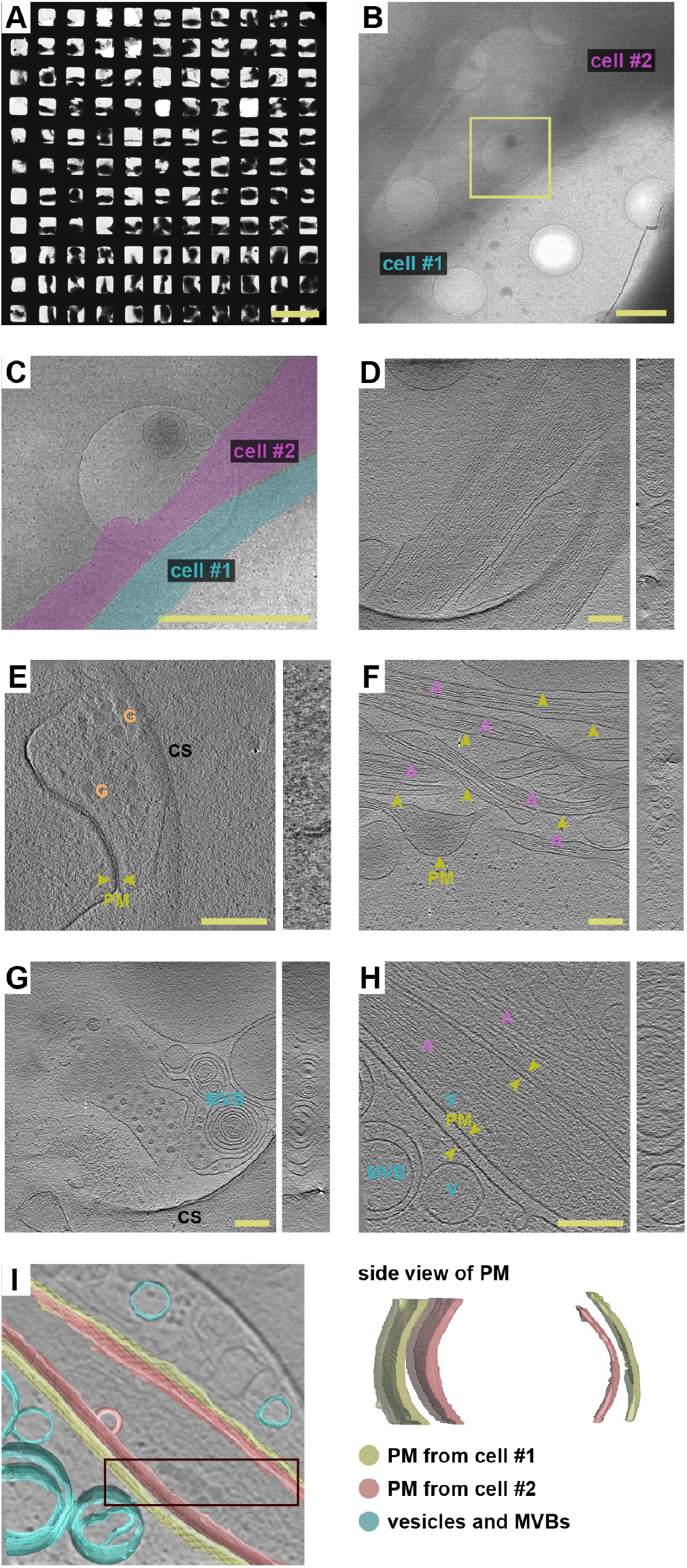
Lattice micropatterning facilitates *in situ* cryo-ET of cell-cell contacts. (A) Representative cryo-TEM grid atlas of ECs adhering to a lattice micropatterned EM grid. Cells on the bottom two rows are oriented perpendicular to the cells in the above rows, as dictated by the micropattern. Such visual cues could aid correlative EM strategies. Scale bar, 200 µm. (B) Magnified cryo-TEM micrograph shows a representative region of interest for the acquisition of a tilt series (yellow box). Scale bar, 2 µm. (C) Zoomed-in image of the region in the yellow box in (B). Contacting cells were colorized for clarity. Scale bar, 2 µm. (D) Tomographic slice of membrane-membrane interaction between cellular protrusions in (C) Scale bar, 200 nm. (E-H) Tomographic slices depict a diversity of membranous structures at cell-cell contacts. *A* actin (purple), *PM* plasma membrane (green), *V* vesicle (cyan), *MVB* multivesicular body (cyan), *G* granule (orange), *CS* carbon support. (D,E) Membrane-membrane interactions between cellular protrusions. (F) Thin, intersecting membrane protrusions. (G) Multivesicular bodies. Scale bars, 200 nm. For full tomograms, see SI. (I) Annotated membranes from (H).

Our reconstructed tomograms revealed a remarkable diversity of filamentous (F) actin-rich membrane protrusions at cell-cell contacts (Fig. 4D-I). We observe several types of endothelial cell-cell contacts consistent with the adherens junctions described by Efimova et al. (13), namely: intercellular bridges with tip-to-tip (Fig. 4E) or lateral (Fig. 4C,D) contacts, and engulfed fingers (Fig. 4H, I). In addition, many tomograms revealed thin, intersecting membrane protrusions between cells, which appear less rigid than expected for filopodia (Fig. 4F). Their function is not immediately clear: They may be thin fibers initiating contact, retraction fibers between parting cells, migrasomes left by retreating cells(14), or suspended protrusions transferring cargo between cells, such as tunneling nanotubes (15), which have recently been structurally analyzed by cryo-ET in neurons (16). Additional investigation using correlative light and electron microscopy techniques will be required to uncover the identity and function of the structures we observed. We also observed a range of vesicle shapes and sizes at endothelial cell-cell contacts (Fig. 4G-I), both within and outside of the plasma membrane.

## Materials and Methods

### Cell culture

We cultured human umbilical vein endothelial cells (ECs, cat. C2591A, Lonza Corporation, Walkdersville, MD) in EGM-2 MV Microvascular Endothelial Cell Growth Medium containing EBM-2 basal medium (Lonza cat. CC3156) and supplemented with penicillin (50 units/mL) and streptomycin (50 µg/ml, Life Technologies, Carlsbad, CA) and the following chemicals from the EGM-2MV BulletKit (Lonza cat. CC-4147): hEGFP, VEGF, hFGF-B, R3-IGF-1, hydrocortisone, and ascorbic acid and 5% FBS. We cultured cells on dishes that were precoated with 0.2% gelatin for *>* 1 h (Sigma-Aldrich, Saint Louis, MO), and maintained them at at 37^*o*^C and 5% CO_2_. We used cells between passages 7 and 12.

### Micropatterning

We plasma treated gold Quantifoil R 2/2 EM grids (657-200-AU, Ted Pella, Redding, CA) for 10 sec at 30 W (PE-50, Plasma Etch, Carson City, NV) and incubated them in a 100 µg/ml solution of poly(l-lysine)-graft-poly(ethylene glycol)(PLL(20)-g[3.5]-PEG(2)) (SuSoS AG, Dübendorf, Switzerland) in phosphate buffered saline (PBS) for 1 h. We rinsed the grids three times in PBS and inserted them carbon-face-down into 25 µL droplets of UV-sensitive photoinitiator (PLPP, Alvéole, Paris, France) contained in custom 4 mm wide silicone wells on a glass-bottom dish. We then placed the dish on the stage of a Leica DMi8 outfitted with an Alvéole PRIMO maskless UV-patterning system and exposed each grid at 2500 mJ/mm^2^ with the lattice micropattern (SI). We tiled the micropattern using the *≈* Leonardo software (22min per 12×12 grid-square region). We rinsed the grids three times in PBS and incubated them in a 0.2% gelatin in BSA bead solution spiked with 10% by volume 1 mg/mL gelatin-Oregon Green 488 conjugate (G13186, Thermo-Fisher Scientific) for 1 h at *≈* 22^*o*^C.

### Cell seeding on micropatterned grids

To obtain the desired distribution of ECs, we suspended cells from an 80% confluent T25 into a 1:8 dilution in cell culture media. We rinsed each grid in PBS, added 10 µL of media to the carbon surface and then added 5 µL of cell suspension at a time until observing ≈ 2 cells/grid-square. After 2 h of incubation, we added 2 mL of warm media to each 35 mm dish and vitrified or fixed the samples after an additional 10-14 h.

### Immunofluorescence and light microscopy

We fixed ECs with 4% paraformaldehyde in PBS for 15 min at *≈* 22^*o*^C. We blocked and permeabilized the ECs in antibody dilution buffer (ADB) made of 0.1% Triton, 1% bovine serum albumin (BSA, Sigma) in PBS for 1 h at *≈* 22 ^*o*^C. We incubated the ECs in a 1:500 primary antibody solution containing Mouse anti-CD144 (VE-Cad, BD Pharmingen Cat. #55561) in ADB overnight at 4 ^*o*^C, and in secondary antibody solution containing Goat anti-mouse IgG Alexa Fluor 647 (Cell Signaling Technology, Cat. #4410) at a 1:1000 dilution in ADB for 1-2 h at *≈* 22^*o*^C. To the latter solution we added Hoechst solution (34580, Thermo-Fisher Scientific) at a 1:1000 dilution to identify nuclei and ActinRed 555 Read Probest reagent (R37112, Thermo-Fisher Scientific) at 1 drop/mL to stain F-actin. Images in Fig. 2A-B were acquired using an inverted Zeiss LSM 780 confocal microscope with a 40X/1.3 NA C-Apo water objective and using Zen Black software (Carl Zeiss). Images used for Fig. 1B, 2C, and S1 were acquired on an inverted Nikon Ti-E microscope equipped with a Heliophor light engine (89 North) and an Andor sCMOS Neo camera using a 20x Plan Apo Lambda air objective lens. Nuclear-junctional distances were measured using ImageJ (S6,S7)(17).

### Atomic force microscopy

We performed atomic force microscopy (AFM) on live and fixed cells cultured on glass bottom dishes using a BioScope Resolve BioAFM in PeakForce QNM mode and a PFQNM-LC-A-CAL cantilever (Bruker, Santa Barbara, CA). We did not perform AFM on cells cultured on EM grids due to the delicate and porous nature of the carbon thin films. There were no discernable differences in cell morphology on micropatterned glass and EM grids (Fig. 3A-B, 2A-B, S2-4). We measured cell-cell contact heights of cells cultured on blanket coated and micropatterned glass bottom dishes with the height sensor channel using Nanoscope Analysis 1.9. The AFM was mounted on a Zeiss Axio Observer Z1 inverted epifluorescent microscope outfitted with a 20x objective.

### Vitrification, cryo-ET, and tomogram reconstruction

We added 3 µL of AuNP solution to each grid (Aurion BSA Gold tracer 10 nm, Fisher Cat. 50-248-12) before vitrifying the grids in a Leica EM GP plunge freezer. We clipped the grids into autogrids and loaded them into an FEI Krios G2 transmission electron microscope (TEM) equipped with an energy filter and a K2 direct electron detector (Gatan). We used SerialEM software (18) to operate the TEM at 300 kV in low-dose mode and acquire tilt series at a magnification of 33,000x (0.4315 nm/px) (Fig 4 D,F,and G) and at a magnification of 64,000x (0.2235 nm/px), using a Volta phase plate to improve image contrast (Fig.4 E and H)(19). Tilt series were acquired with 2^*o*^ steps between -60^*o*^ and +60^*o*^ and a defocus range between -2 µm and -5 µm. We motion corrected images with MotionCor2 software (20), aligned them using gold fiducials, and reconstructed tomograms using the IMOD software package, version 4.9.1 (21). 3D reconstructions were calculated using the weighted-back projection using IMOD (21). We segmented the tomogram volume in Fig. 4I using Amira (FEI/Thermo Fisher Scientific) software.

## Supporting information

Digital micropattern file

Tomogram from Figure 4E

Tomogram from Figure 4F

Tomogram from Figure 4H

Tomogram from Figure 4G

## Acknowledgements

We thank Dunn and Weis lab members for helpful discussion and M. Azubel, R.G. Held, and J. Leitz for helpful advice. The project described was supported, in part, by Award Number 1S10OD021514-01 from the National Center for Research Resources (NCRR). Its contents are solely the responsibility of the authors and do not necessarily represent the official views of the NCRR or the NIH. Work was performed in part in the nano@Stanford labs, which are supported by the National Science Foundation as part of the National Nanotechnology Coordinated Infrastructure under award ECCS-1542152. Fig 1A was created with BioRender.com. This work was supported by NIH F32GM125113 (CGV), R01HL128779 and R35GM130332 (ARD), R01GM119948 and R35GM131747 (WIW), and an HHMI Faculty Scholars Award (ARD).

## Author Contributions

L.E., C.G.V., W.I.W., and A.R.D. designed research; L.E., C.G.V., E.A.M., B.M.S. and M.P.W. performed research and analyzed data; L.E. and C.G.V. wrote the paper; A.R.D and WIW edited the paper.

## Supporting Information (SI)

### Supporting material

Several figures that support the results reported in the manuscript.

### Digital micropattern file

The lattice micropattern .TIF file is compatible with the Alveole Primo micropatterning system.

### SI movies

Full tomograms for figures 4D-H.

## Notes

### Competing Interest Statement

The authors have declared no competing interest.

