## Supplementary figures and images for "Lattice micropatterning for cryo-electron tomography studies of cell-cell contacts"

### Digital micropattern file

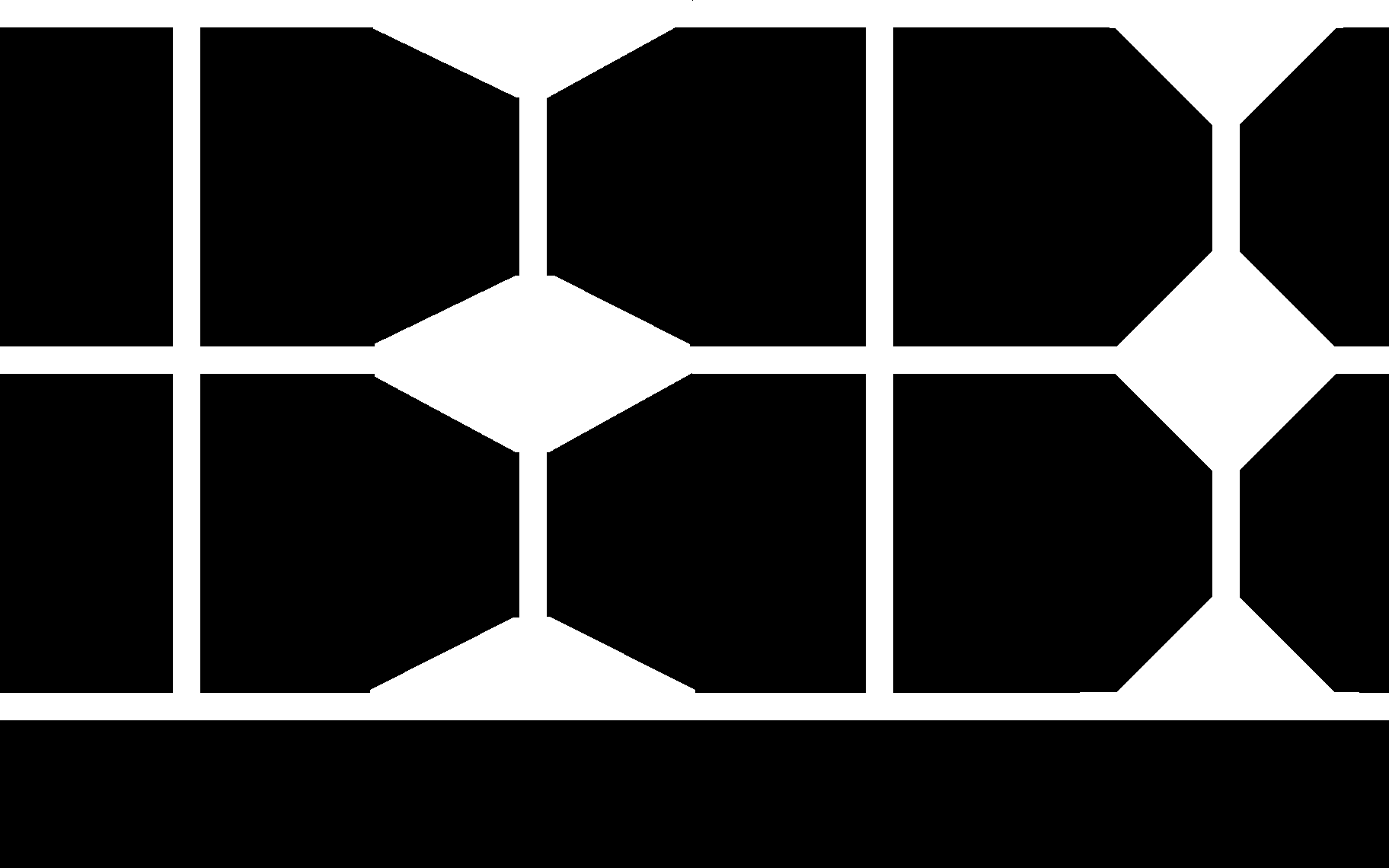
