## Supplemental figures for "Lattice micropatterning for cryo-electron tomography studies of cell-cell contacts"

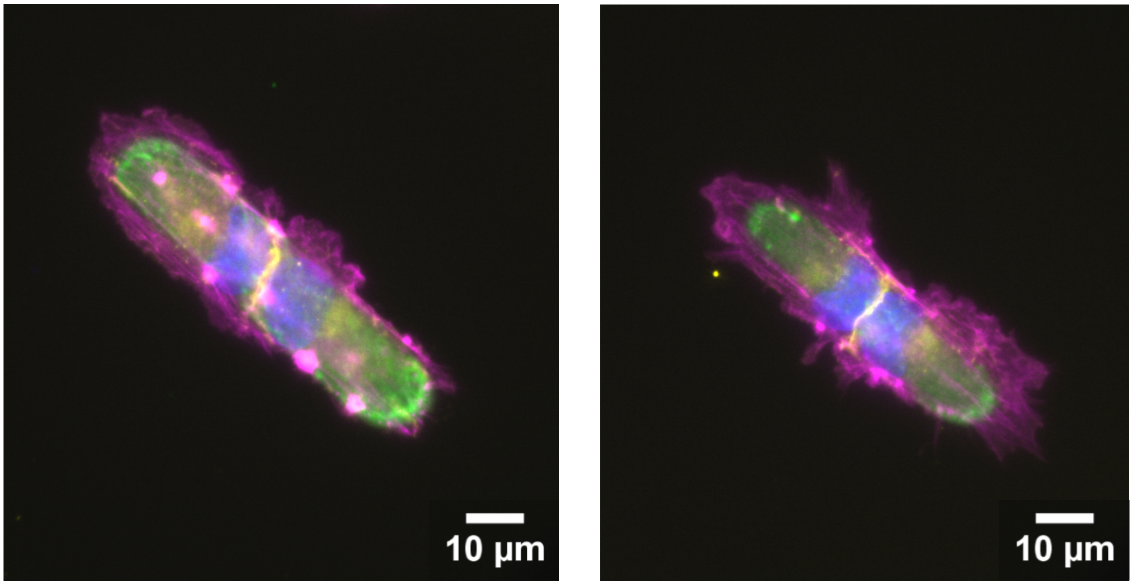


**Fig. S1:** Endothelial cell (EC) nuclei in cell pairs assembled on micropatterned islands position nuclei in close proximity to cell-cell contacts. Fluorescent micrographs depict cell-cell contacts (VE-cadherin, yellow), nuclei (DAPI, blue), actin (magenta) in cell doublets attached to gelatin micropatterns (green). Left micropattern: 120 x 20 μm. Right micropattern: 80 x 10 μm.

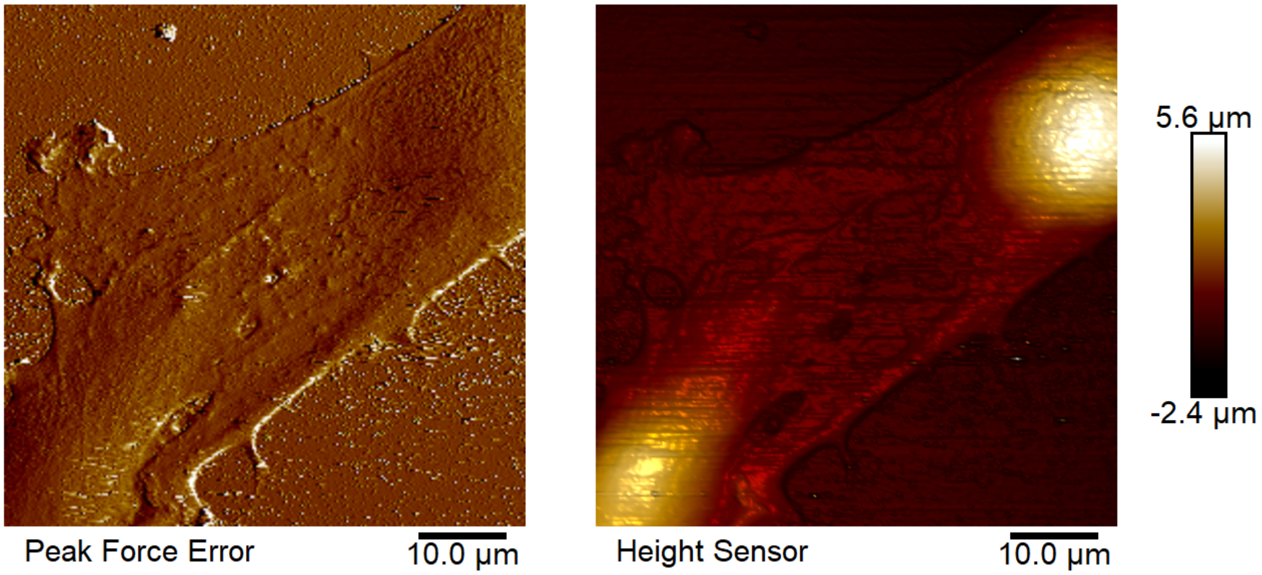


**Fig. S2**: <u>EC nuclei are micron scale in height.</u> Live cell atomic force microscopy (AFM) of endothelial cell-cell contact 12 h after cells were seeded on gelatin-coated glass. Left: peak force error channel. Right: height sensor channel.

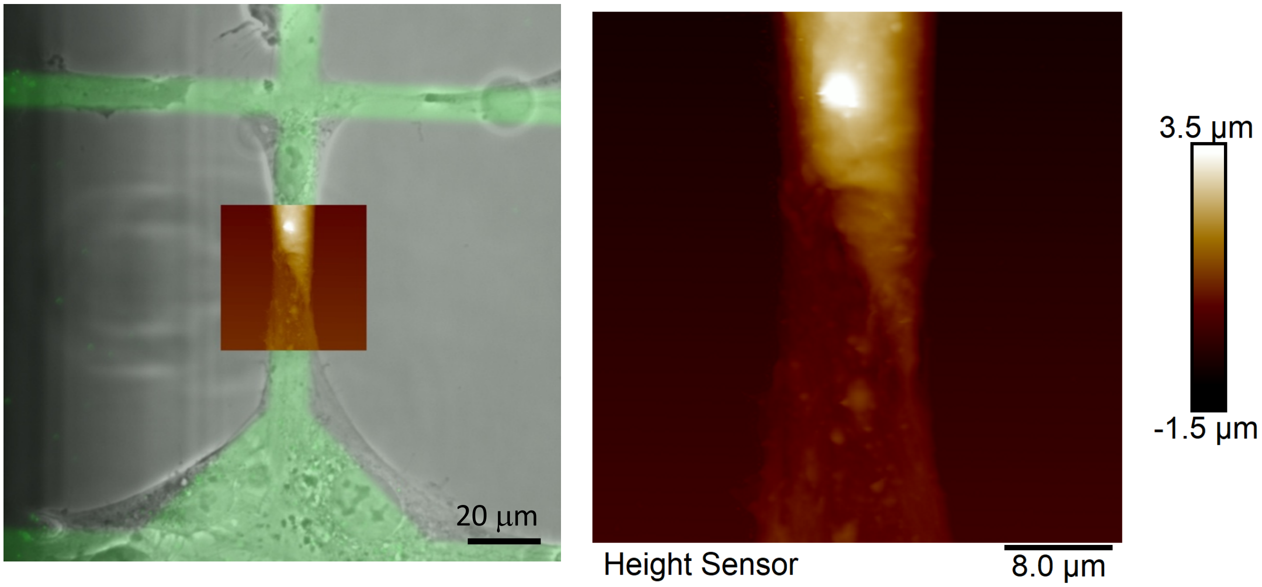


**Fig. S3**: <u>Proximity to cell nuclei increases cell-cell contact thickness.</u> Left: Brightfield image of ECs on micropatterned glass (grayscale) overlaid with fluorescence image of gelatin micropattern (green). Inset, magnified at right, shows the height sensor channel of an AFM scan an a cell-cell contact. The upper cell is taller than the lower cell at the cell-cell contact due to its proximity to the cell nucleus positioned above it.

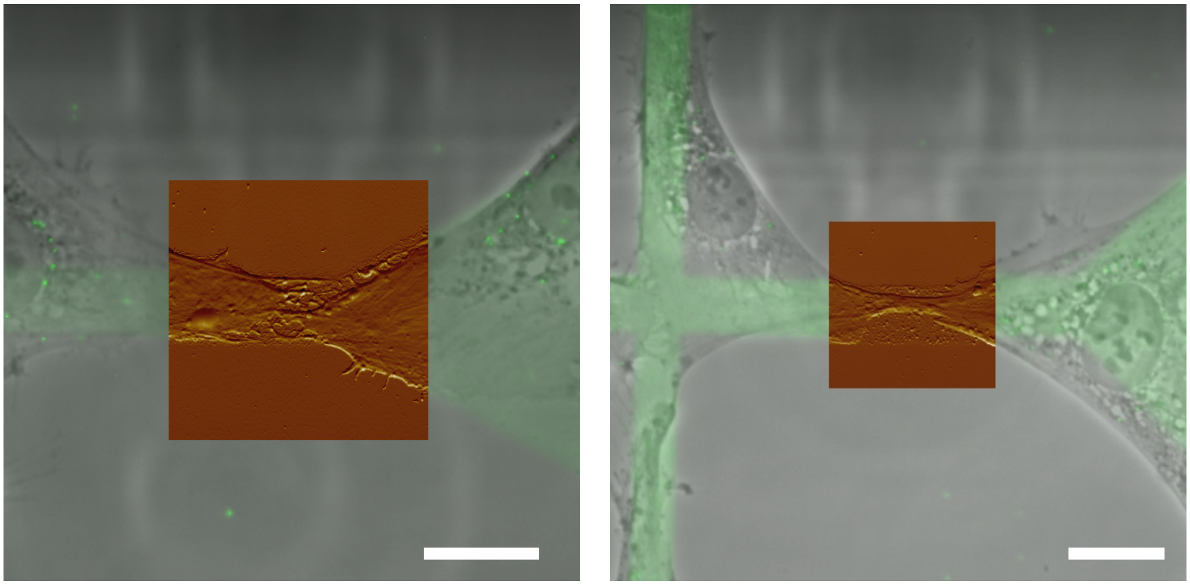

**Fig. S4**: ECs use ECM tracks as templates for cell-cell contact assembly and/or cell retraction. Left: Brightfield image of contacting ECs on micropatterned glass (grayscale) overlaid with fluorescence image of gelatin micropattern (green). Inset shows the topography of thin protrusions where cells are in contact via the AFM peak force error channel. Scale bars, 20 $\mu$m.

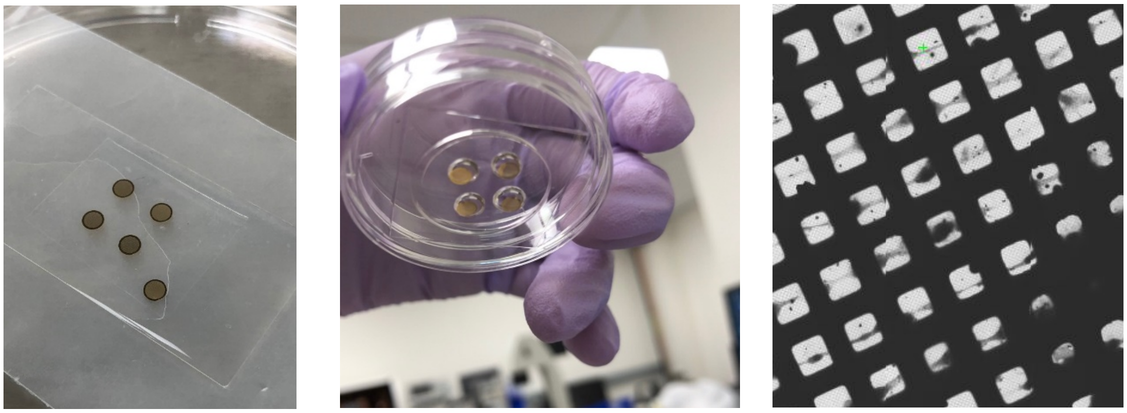

**Fig. S5**: Grid handling steps for maintaining grid integrity during maskless photopatterning. (A) EM grids are mounted carbon-face-up on silicone sheets on glass slides for plasma treatment. They are passivated by incubation in PLL-PEG on parafilm. (B) Custom silicone stencils are used to create wells 4 mm in diameter that can contain liquid droplets during photopatterning or protein incubation steps. Grids are placed carbon-face down in PLPP droplets for photopatterning. (C) Cryo-transmission electron micrograph shows ECs adhering to a lattice micropatterned EM grid with intact holey carbon film.

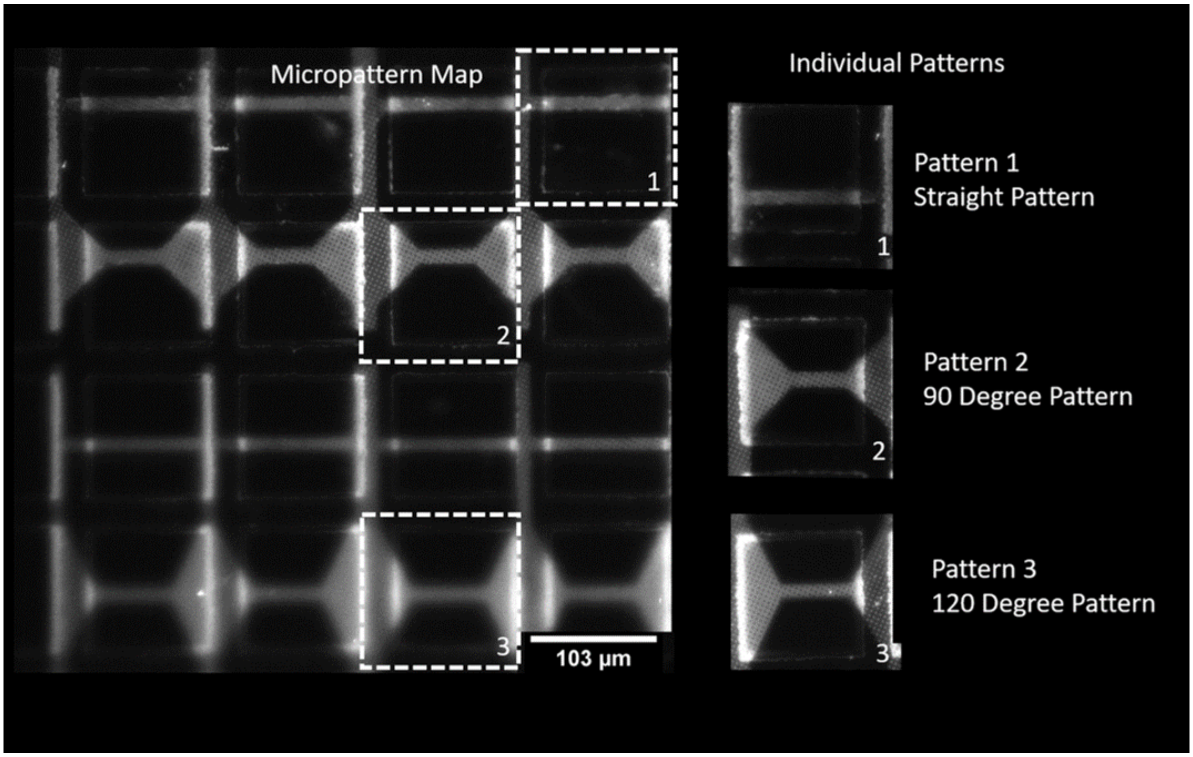


**Fig. S6**: <u>ECM micropatterns tested.</u> EM grid with lattice micropattern (left) features three different patterns (right). Thin horizontal and vertical tracks are 10 µm wide.

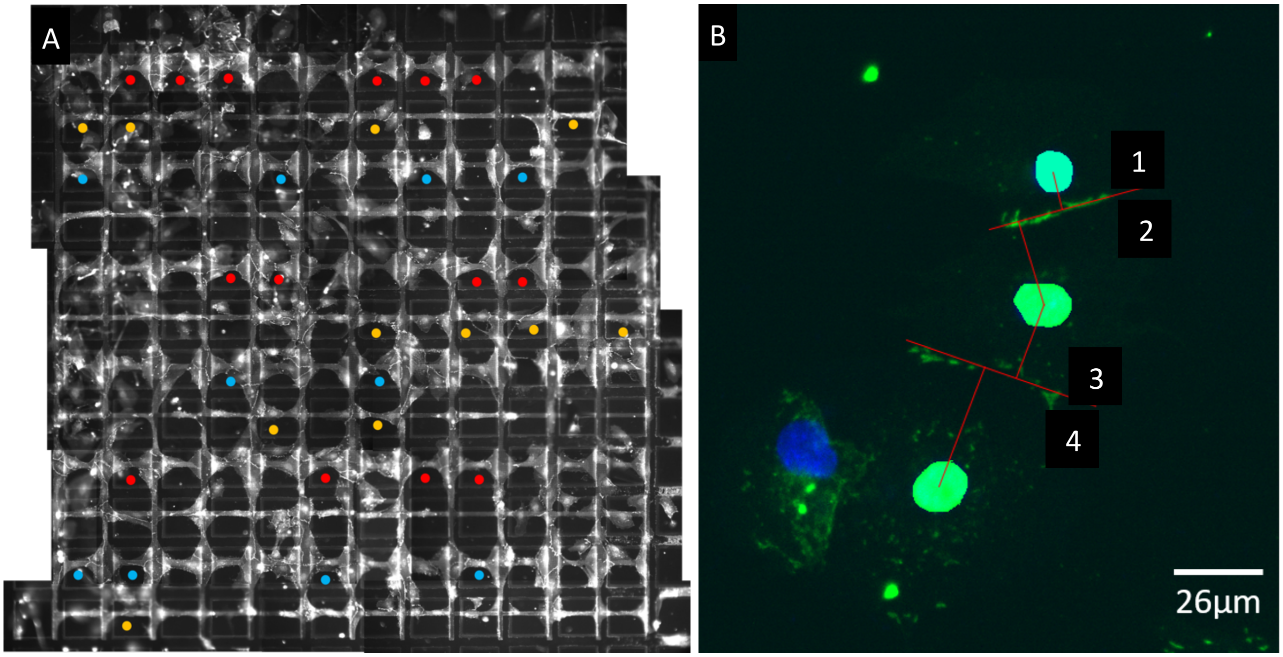


**Fig. S7**: <u>Analysis of cell assembly on micropatterned EM grids</u>. (A) Representative fluorescent micrograph of fixed ECs stained for VE-cadherin (white) on micropatterned EM grids. Circles indicate cell-cell contacts where nuclear-junctional distances were measured. Each color corresponds to a different micropattern type: straight (yellow), 90° (red), 120° (blue). (B) Example of nuclear-junctional measurements. Nuclei are segmented, VE-cadherin is green.
